# Evaluating and Classifying Gentleness in VR-Based Surgical Simulation: A VR+fNIRS Study

**DOI:** 10.64898/2026.01.02.697327

**Authors:** Suveyda Sanli, Hasan Onur Keles

## Abstract

Gentleness, defined as the ability to handle tissues delicately and minimize unnecessary force, is a key indicator of surgical proficiency. Objective and real-time assessment of gentleness in virtual reality (VR)–based training can enhance the understanding of both psychomotor and cognitive aspects of surgical skill. This study evaluates and classifies participants’ gentleness during VR-based surgical simulations using fNIRS-derived hemodynamic features. We trained and compared several machine learning models to assess performance. Twenty-three volunteers with no prior laparoscopic experience performed a virtual reality–based laparoscopic double-grasper task while hemodynamic activity over frontal and motor cortical areas was recorded using eighteen fNIRS channels. Alongside fNIRS, we collected subjective workload (NASA-TLX), error numbers, and a VR gentleness score. This task involves using two grasper tools simultaneously to perform the tissue like balloon manipulation in a VR environment. We extracted temporal features (slope, RMS, standard deviation) and trained machine learning models to classify performance levels based on cortical activation. Labels were binarized as low vs. high using median splits for the gentleness score. Models were evaluated with stratified 5-fold cross-validation and summarized by accuracy. Results showed stronger right-frontal HbO activity and increased left-motor HbR responses in the low-performance group, suggesting greater cognitive effort and less efficient motor strategies during VR-based laparoscopic manipulation. Across classifiers and feature sets, slope-based features consistently outperformed variability- and amplitude-based metrics. Among the tested models, HbR slope features achieved the best overall classification performance, with the highest accuracy obtained using K-Nearest Neighbor and Random Forest classifiers (accuracy ≈ 0.89, AUC up to 0.97). These findings demonstrate that fNIRS-derived hemodynamic dynamics can reliably discriminate between high and low VR performance levels, supporting their potential use in automated performance assessment and neuroadaptive feedback frameworks for VR-based surgical training.

## INTRODUCTION

Virtual reality (VR) simulators have become increasingly integrated into medical education(1). Over the past decade, their use has expanded in the field of surgical training especially and VR platforms continue to maintain strong popularity as training tools(2,3). These platforms are now routinely implemented for basic psychomotor skill acquisition, procedural rehearsal, team-based simulation, and competency-based assessment. However, their overall effectiveness and the mechanisms through which they enhance surgical performance remain subjects of ongoing discussion(4–6). Specially, the underlying neural and cognitive mechanisms remain to be fully understood(7). Combining the VR headsets with neuroimaging techniques offers a important approach to uncovering the neural and cognitive mechanisms underlying surgical skill acquisition and performance during VR-based training(8–10).

Surgical training has traditionally measured success with skills like speed, accuracy, and error rates. However, an equally vital skill, namely gentleness, is often overlooked in these assessments(11). This study focuses specifically on this critical skill. VR simulators present a valuable tool for assessing essential surgical competencies like gentleness, which is a fundamental indicator of competence across all surgical domains. As excessive force can cause tissue damage and prolong patient recovery. Importantly, surgical gentleness encompasses more than motor execution alone; it involves sustained attention, anticipatory planning, inhibition of unnecessary force, and precise regulation of movement under limited sensory feedback(12–14). Because of its multidimensional nature, numerous surgical education frameworks have emphasized the need to objectively quantify gentleness and incorporate it into modern assessment tools. Gentleness has been incorporated into procedural assessment frameworks, particularly in minimally invasive surgery, where excessive force may lead to unintended tissue damage, postoperative complications, and prolonged recovery times. Evidence shows that surgical skill is a strong predictor of patient outcomes, and gentleness is one of the primary contributors to overall surgical competency(15).

The Fundamentals of Laparoscopic Training (FLS) curriculum is a mandatory component of laparoscopic education and must be completed prior to board certification in some countries(16). Safe surgery has been defined by principles such as gentle handling of tissues, meticulous hemostasis, avoidance of dead space, and adherence to precise surgical technique. Among these principles, gentleness has increasingly gained recognition as a measurable and essential aspect of operative performance assessment.

Despite its importance, surgical skill evaluation traditionally relies on subjective assessments within an apprenticeship-model training paradigm(16,17). Such evaluations lack objective structure and fail to provide meaningful, real-time feedback to trainees(18). In contrast, proficiency-based simulation training has demonstrated superior outcomes, leading to fewer errors and complications in the operating room and improving trainee performance across multiple metrics. As a result, many surgical training programs have adopted simulation-based curricula to support skill acquisition and assessment(19). Given the growing emphasis on competency-based education and the clear clinical relevance of gentleness, there is a need for objective and scalable tools that quantify this skill(11). Addressing this gap is the important of the present study. There is a growing need to utilize VR simulators in combination with functional neuroimaging modalities to investigate the neural mechanisms underlying motor learning, cognitive workload, and decision-making during simulated surgery

In minimally invasive surgery, appropriate manipulation forces, bimanual coordination, and gentleness toward soft tissue are essential to avoid unintended tissue damage and to accomplish stable and precise handling(17,20–22). Traditional metrics such as error counts or task duration cannot fully capture these subtle aspects of motor behavior. Therefore, assessing gentleness, defined as the ability to perform precise manipulation with minimal unnecessary force, has become a parameter in objective skill quantification.

The double graspers manipulation task, which requires users to transfer a soft, deformable object (balloon) using a pair of grasper tools, provides a natural environment for quantifying gentleness. Successful performance demands bilateral coordination, adequate pressure control, and strategic motor planning to avoid tearing, dropping, or excessively deforming the object. Consequently, the performance score obtained from this task is considered a sensitive indicator of soft-tissue handling skill, and can differentiate novice behavior from more controlled manipulation strategies(11,23).

Functional near-infrared spectroscopy (fNIRS) provides a portable and ecologically valid functional neuroimaging method to investigate cortical hemodynamics during physically interactive VR-based training(24–27). Previous studies have shown that prefrontal activation reflects cognitive load, motor planning, and adaptive processes during demanding motor tasks, whereas sensorimotor responses are modulated by task difficulty and motor proficiency(16,28–31). By integrating VR performance metrics with fNIRS-derived cortical activation patterns, it becomes possible not only to assess how well participants perform, but also to characterize how they allocate cognitive and motor resources to achieve gentle manipulation. Machine learning approaches have been increasingly applied to predict surgical skill levels using neuroimaging data(32–35); however, there is limited research exploring their use in VR-based surgical training environments. Existing studies often rely on relatively simple models, non-neuroimaging metrics, and lack systematic comparisons across different machine learning algorithms(36,37).

In this paper, we investigate the cerebral hemodynamic prediction of gentleness by dividing VR gentleness performance scores into two groups (high vs. low). Classification is performed using both prefrontal and motor cortical activation features, demonstrating that optical signals contain reliable information to discriminate levels of gentleness during VR-based surgical training. In addition, we examine the relationship between subjective workload and gentleness, showing that higher gentleness is associated with reduced perceived workload.

The proposed methodology highlights that combining fNIRS imaging with machine learning provides an objective and skill-relevant metric that complements subjective assessments in VR environments. This neuroimaging-based framework offers quantitative and standardized measures that can support professional certification and surgical education during VR training. This is the first study conducted with this participant group, and future work will expand the study to include expert and novice surgeons using the insights gained from this initial phase.

## METHODS

### Materials and Methods

#### Participants

The study involved 23 volunteer participants with no prior laparoscopic experience. The participants had an average age of 27 ± 6 years. All participants are right-handed. All participants provided written informed consent prior to the experimental session, in accordance with the Declaration of Helsinki. The study protocol was reviewed and approved by the Ankara University Human Research Ethics Committee (Approval No: 2024000333-1), and data were collected between 20/06/2024 and 20/08/2025.

#### Experimental Design

Participants performed the Laparoscopic Double Grasper Task in a VR environment while fNIRS data were recorded in a single session. Prior to data collection, a brief 3-minute practice trial was conducted to familiarize participants with the virtual setting. To evaluate surgical gentleness and precision, a 3D virtual reality simulator (the Laparoscopic Double Grasper Task) was developed in Unity 3D and deployed on the Meta Quest 3 headset(11). The wireless setup allowed natural interaction and real-time tracking of instrument movements, while continuous visual feedback enhanced spatial awareness and depth perception throughout the task. Participants first completed a 2-minute resting-state baseline while keeping their eyes open and minimizing movement, which served as a reference for task-related activity. They then performed the 5-minute Laparoscopic Double Grasper Task, manipulating a soft, tissue-like balloon with two virtual graspers in the VR environment. The system automatically calculated performance scores based on accurate transfers and placements, while errors (balloon bursts due to excessive pressure) were manually noted.

#### Laparoscopic Double Grasper Task

The Laparoscopic Double Grasper Task was designed to evaluate participants’ coordination and gentle tissue handling in a simulated laparoscopic environment. The main objective of the double graspers task was to use both hands to transfer the balloon, a soft body, from the right box side of the scene into the left box via the use of haptic devices. Balloon stiffness was set to 0.1 again to correctly simulate soft tissue like behavior. In the virtual operating room , two boxes were placed on a table and separated by a central frame. A soft, balloon-shaped object representing biological tissue was initially positioned in the left box.

Participants used the left grasper to pick up the balloon, gently transferred it through the central opening, and placed it in the right box with the right grasper. They then repeated the motion in reverse to complete one full transfer cycle. The balloon’s stiffness was adjusted to mimic fragile tissue, requiring precise control. Excessive pressure caused the balloon to burst with an audible pop, which was recorded as an error. If the grasping force was insufficient, the balloon slipped away, causing time loss within the five-minute task period. A real-time score display in the VR scene provided performance feedback. Transfers performed near the frame’s center earned up to 50 points, and successful placements added 25 points. The final score reflected the total number and accuracy of completed transfers. The visual components of the VR environment are shown in Figure 2.

#### Data Collection

We used a wearable continuous wave fNIRS device (Brite, Artinis Medical Systems, The Netherlands). The device is a continuous-wave system using two wavelengths (760 nm and 850 nm) to record back-reflected light intensity. The fNIRS system comprised 10 sources and 8 detectors arranged in 2×4 and 2×5 probe layouts over the frontal and motor cortical regions, respectively (Figure 1). The resulting 18 channels were symmetrically divided, with 9 channels placed over each hemisphere. Each hemisphere was monitored by 5 prefrontal (frontal) channels and 4 motor cortex channels (Figure 1). Optode–scalp contact and signal quality were rigorously verified prior to beginning the recording session. Sources and detectors are separated by approximately 30 mm (i.e., long-separation channels). During the fnirs recording, the number of error were also recorded. All behavioral and fNIRS signals were synchronized to ensure precise temporal alignment. After the task, participants completed the NASA-TLX survey to rate perceived workload across mental, physical, temporal demand, performance, effort, and frustration, each on a 20-point visual analog scale.

**Figure 1.**
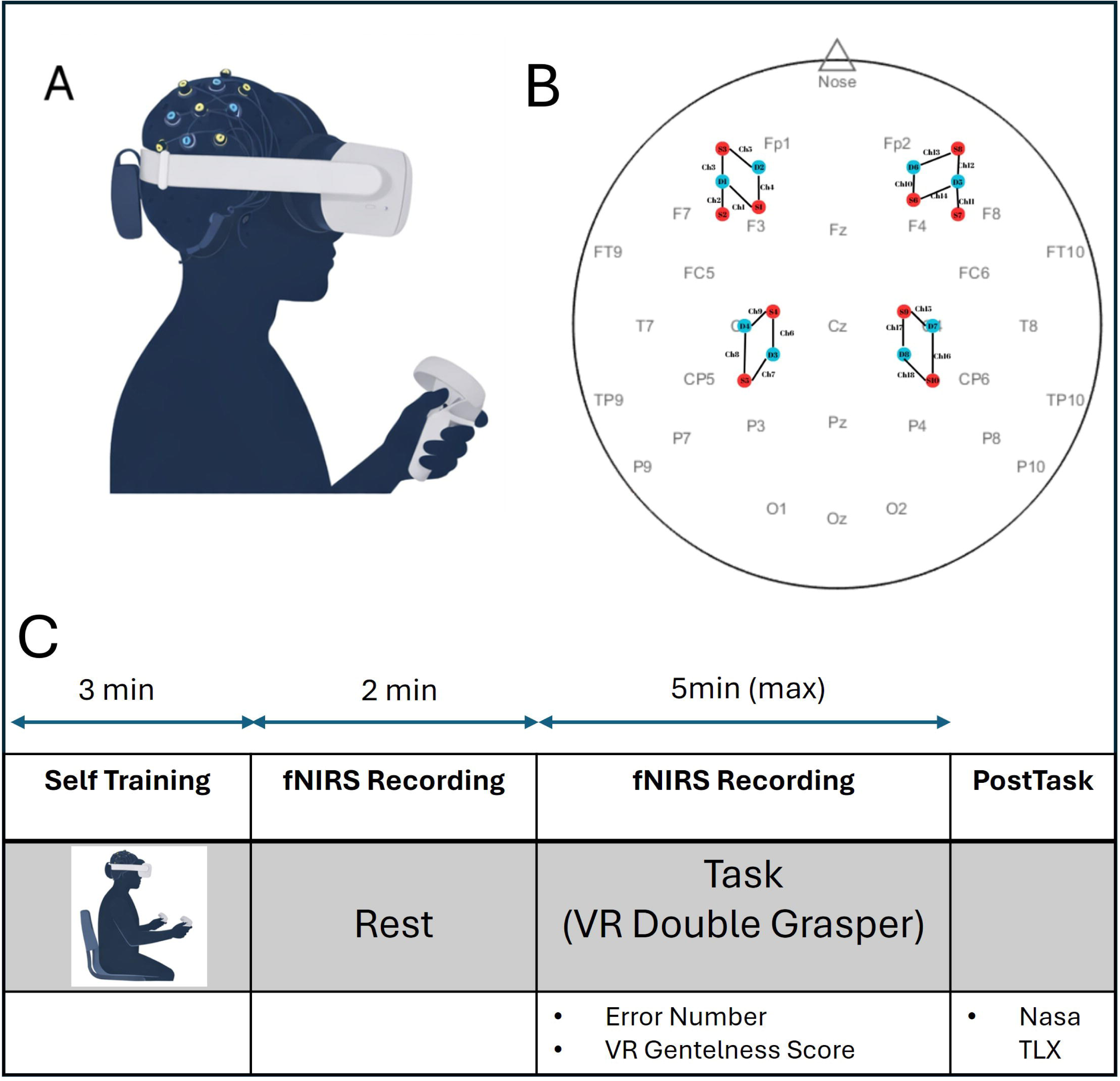
VR double-grasp task used in laparoscopic simulation. The VR environment requires users to grasp and transfer virtual objects (ball) between two target areas using two laparoscopic controllers. Performance is scored automatically in real time based on task completion and errors (e.g., drops or missed grasps), and the cumulative score is displayed on the screen

**Figure 2.**
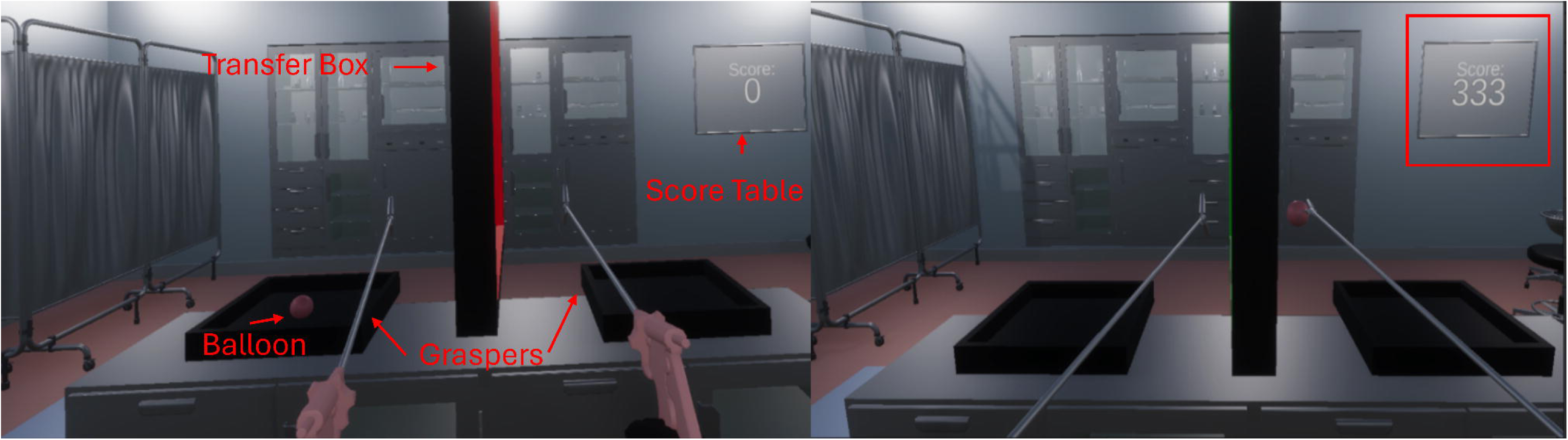
Experimental setup and fNIRS sensor configuration during VR-based laparoscopic training. **(A)** A participant wearing the fNIRS cap and VR headset while performing the laparoscopic simulation task using a handheld controller. **(B)** Optode layout showing the placement of sources and detectors over frontal and motor cortices following the international EEG 10–20 system.**(C)** Timeline of the experiment illustrating the calibration period, VR task execution, and recording duration during free laparoscopic training.

### Data Analysis

#### fNIRS signal preprocessing

Homer3 and MATLAB (The MathWorks, Inc., Natick, MA, USA) were used for the preprocessing and analysis of the fNIRS data. Channels with poor signal quality were identified based on raw light intensity amplitude thresholds outside the acceptable range (1.7772 × 10⁻□= to 1.9 arbitrary units). Following channel pruning, the signal was converted into optical density (OD). After a visual inspection of the data, wavelet-based method was applied to correct for motion artifacts(38). To remove high-frequency oscillations and physiological noise, the corrected OD signal was processed using a sixth-order Butterworth low-pass filter with a frequency range of 0.01–0.5 Hz. Finally, the changes in optical density were converted into concentration changes of oxygenated hemoglobin (HbO) and deoxygenated hemoglobin (HbR) by employing the Modified Beer-Lambert Law(39).

To assess localized cortical activity, we calculated the standard deviation of oxyhemoglobin changes within non-overlapping 10-second windows for each measurement channel. Since a stronger evoked hemodynamic response typically increases the signal variability within a given window, we used these standard deviation values as an indicator of prefrontal activation. This method is preferable to using the window mean in certain cases, such as when the evoked response is short-lived and followed by a subsequent drop in the signal(16).

We also applied this windowed standard deviation to identify significant motion artifacts, specifically those with amplitudes exceeding those caused by normal physiological activity. A smaller standard deviation would indicate a less severe artifact. For each channel, we computed the standard deviation across all 10-second windows and then determined the median absolute deviation (MAD) of those values. Any window whose standard deviation fell more than 4.5 MAD units above the median was labeled as an outlier and removed from further analysis.

#### Feature extraction and machine learning classification

From the preprocessed HbO and HbR signals, several statistical features were computed for 5 minutes task period, including the *standard deviation*, *root mean square (RMS)*, and *slope* values. Feature extraction was performed within 10-second windows, and each window was treated as an independent data sample. To investigate whether fNIRS-derived features could discriminate between different VR performance scores were divided into *low* and *high* groups and used as binary class labels. All features were normalized using *StandardScaler*, and model evaluation employed 5-fold cross-validation with a 20% test split. Several machine learning classifiers were trained and compared, including Random Forest, Support Vector Machine (SVM) and K-Nearest Neighbors (KNN). GridSearchCV was used for hyperparameter optimization. Model performance was evaluated using accuracy, F1-score, precision, and area under the ROC curve (AUC), and results were visualized using confusion matrices. Figure 3 summarizes the experimental workflow from raw fNIRS data acquisition to performance prediction

**Figure 3.**
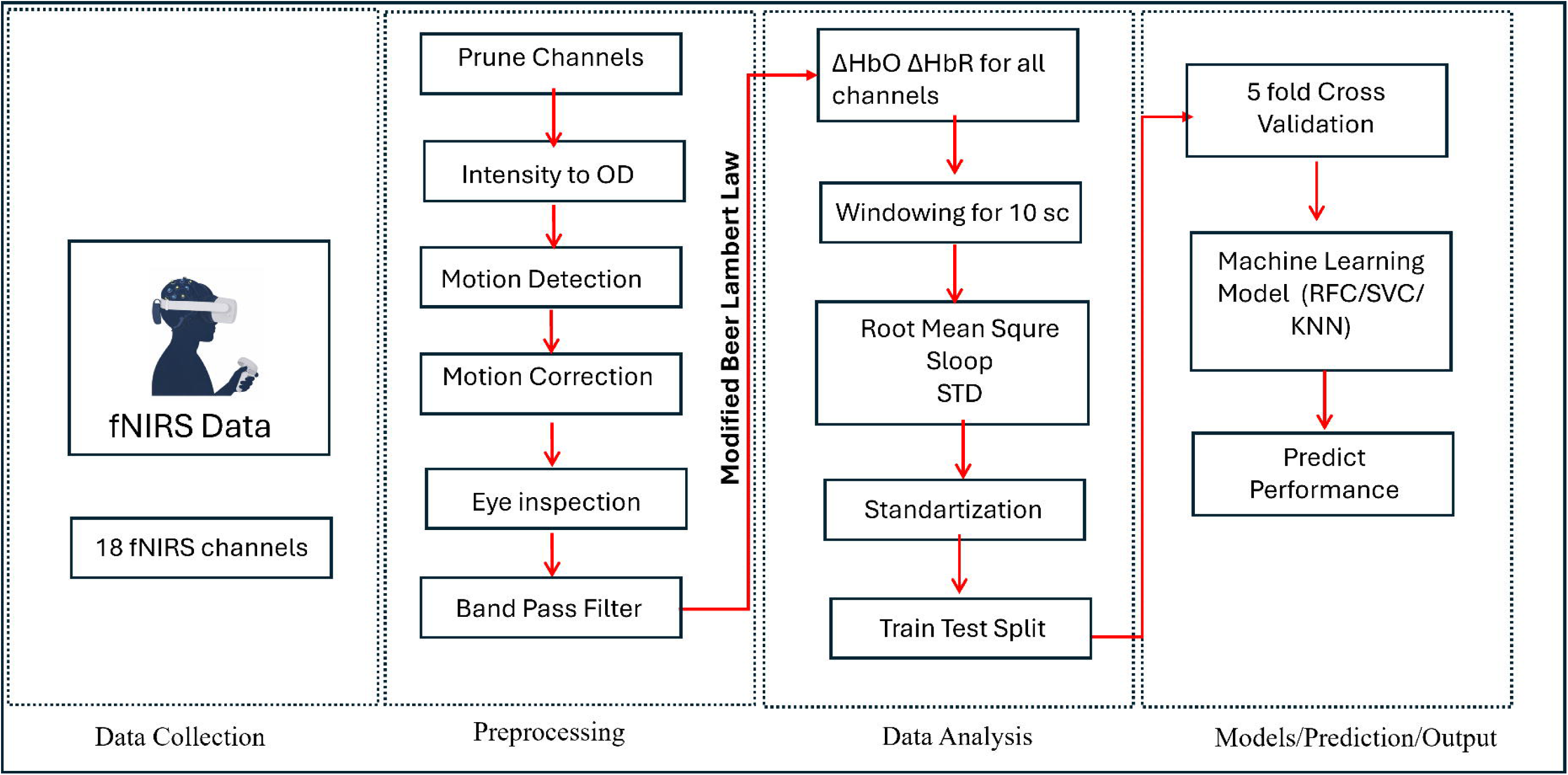
The workflow for fNIRS data acquisition, preprocessing, and machine learning classification.

#### NASA-TLX and VR score evaluation

To explore the relationship between subjective workload and objective task performance, Pearson correlation analyses were conducted between each NASA-TLX subscale and VR performance score. Subscales included *mental demand, physical demand, temporal demand, performance, effort,* and *frustration*, as well as the *total workload score*. Correlation strength and direction were quantified using *r* values, with statistical significance set at *p* < 0.05.

### Statistical analysis

When conducting a regression analysis comparing 2 numerical variables, linear fit with analysis of variance was used. The descriptive results comparing two groups, such as completion time v game experience; completion time v laparoscopy experience; NASA total v game experience and NASA total v laparoscopy experience, HbO changes contained non-paired data. In order to assess the statistical significance of the difference between two groups of non-paired results, we used the non-parametric Kolmogorov test . We did not utilize null-hypotheses whose rejection would have required corrections for multiple comparisons or false discovery.

## RESULTS

We report behavioral, subjective, and neurophysiological results from a VR-based laparoscopic training task involving 23 participants. Figure 5 shows the relationship between the gentleness performance score and perceived workload (NASA-TLX) and subscales. Figure 6 presents the hemodynamic differences between high and low gentleness groups across prefrontal and motor regions. Table 1 summarizes the machine-learning classification of gentleness using fNIRS features extracted from cortical signals, specifically SLOPE, RMS, and STD. Figure 7 displays the corresponding confusion matrix, highlighting the predictive performance of the proposed model.

**Table 1.** Performance Metrics of Machine Learning Models Classified by fNIRS Features.

| Features | Classifier | Accuracy | F1 Score | Precision | Recall | AUC Score |
| --- | --- | --- | --- | --- | --- | --- |
| HbO SLOPE | RFC | 0.7692 | 0.7724 | 0.8000 | 0.7467 | 0.8834 |
|  | SVC | 0.8601 | 0.8718 | 0.8395 | 0.9067 | 0.9071 |
|  | KNN | 0.8392 | 0.8535 | 0.8171 | 0.8933 | 0.9134 |
| HbR SLOPE | RFC | 0.8881 | 0.8904 | 0.9155 | 0.8667 | 0.9656 |
|  | SVC | 0.8252 | 0.8428 | 0.7976 | 0.8933 | 0.9157 |
|  | KNN | 0.8881 | 0.8961 | 0.8734 | 0.9200 | 0.9325 |
| HbO STD | RFC | 0.7343 | 0.7397 | 0.7606 | 0.7200 | 0.8084 |
|  | SVC | 0.7133 | 0.7285 | 0.7237 | 0.7333 | 0.7888 |
|  | KNN | 0.7692 | 0.7975 | 0.7386 | 0.8667 | 0.8508 |
| HbR STD | RFC | 0.6713 | 0.6466 | 0.7414 | 0.5733 | 0.7969 |
|  | SVC | 0.6573 | 0.6667 | 0.6806 | 0.6533 | 0.7871 |
|  | KNN | 0.8462 | 0.8514 | 0.8630 | 0.8400 | 0.9196 |
| HbO RMS | RFC | 0.7622 | 0.7671 | 0.7887 | 0.7467 | 0.8173 |
|  | SVC | 0.8042 | 0.8228 | 0.7831 | 0.8667 | 0.8780 |
|  | KNN | 0.8042 | 0.8182 | 0.7975 | 0.8400 | 0.8698 |
| HbR RMS | RFC | 0.6783 | 0.6515 | 0.7544 | 0.5733 | 0.7971 |
|  | SVC | 0.7343 | 0.7500 | 0.7403 | 0.7600 | 0.8257 |
|  | KNN | 0.7902 | 0.7917 | 0.8261 | 0.7600 | 0.8790 |

A total of 23 participants were included in the VR performance analysis. Figure 4A shows that the distribution of total VR scores exhibited wide variability, ranging from 201 to 2,163. Based on the median value (Median = 815), participants were divided into two performance groups: LOW VR (≤ 815; n = 12) and HIGH VR (> 815; n = 11). This median-based split was used for subsequent group-level comparisons.

**Figure 4.**
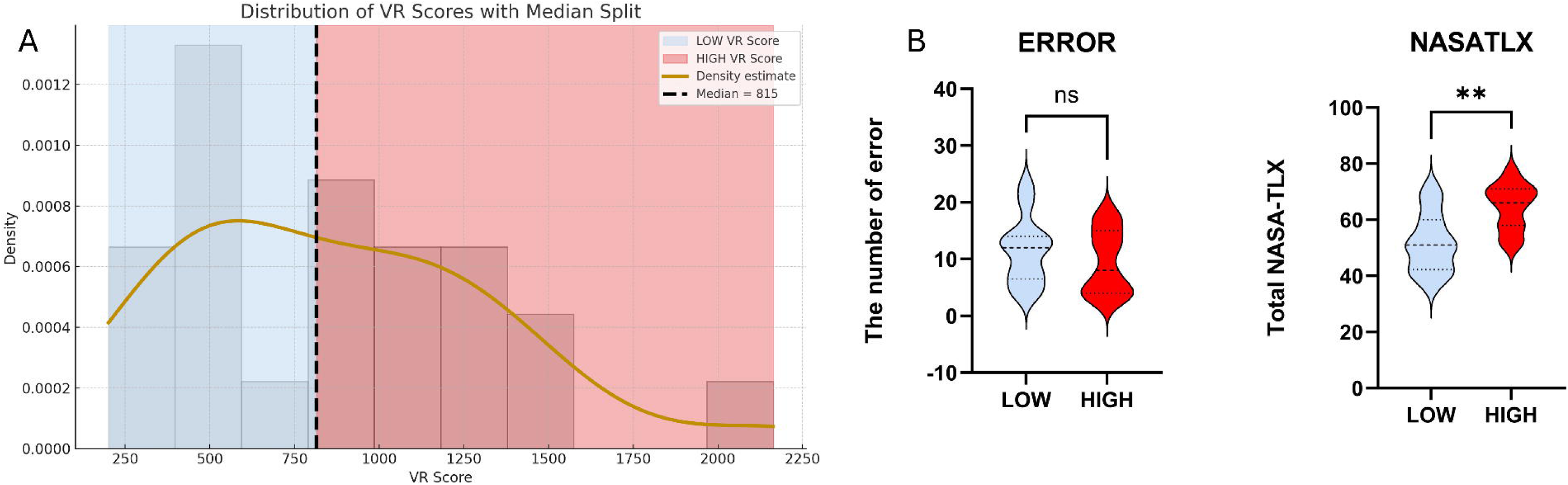
(A) Histogram of total VR scores (n = 23) showing a wide variability. Participants were divided into LOW and HIGH VR performance groups based on the median value (median = 815). (B) Comparison of task performance metrics between LOW and HIGH groups. The number of errors did not differ significantly between groups (ns), whereas the HIGH VR group reported significantly higher overall workload on the NASA-TLX (p < 0.01).

Figure 4B shows that the HIGH VR score group did not exhibit a significant reduction in the number of errors compared to the LOW VR score group (ns; p > 0.05), indicating that higher VR performance was not necessarily associated with fewer mistakes. However, the HIGH VR group reported significantly higher NASA-TLX workload scores (p < 0.01), suggesting that participants who performed better experienced greater perceived cognitive and physical demand during the task.

Figure 5 shows that Regression analyses between the VR Gentleness Score and NASA-TLX subscales showed significant relationships for Physical Demand (p = 0.05) and Performance (p = 0.005). Trends in the same direction were observed for Mental Demand, Temporal Demand, and Effort, although these did not reach significance. Frustration showed a weak negative relationship with the gentleness score. The analysis of the total NASA-TLX score also revealed a significant association with the VR Gentleness Score (p = 0.01).

**Figure 5.**
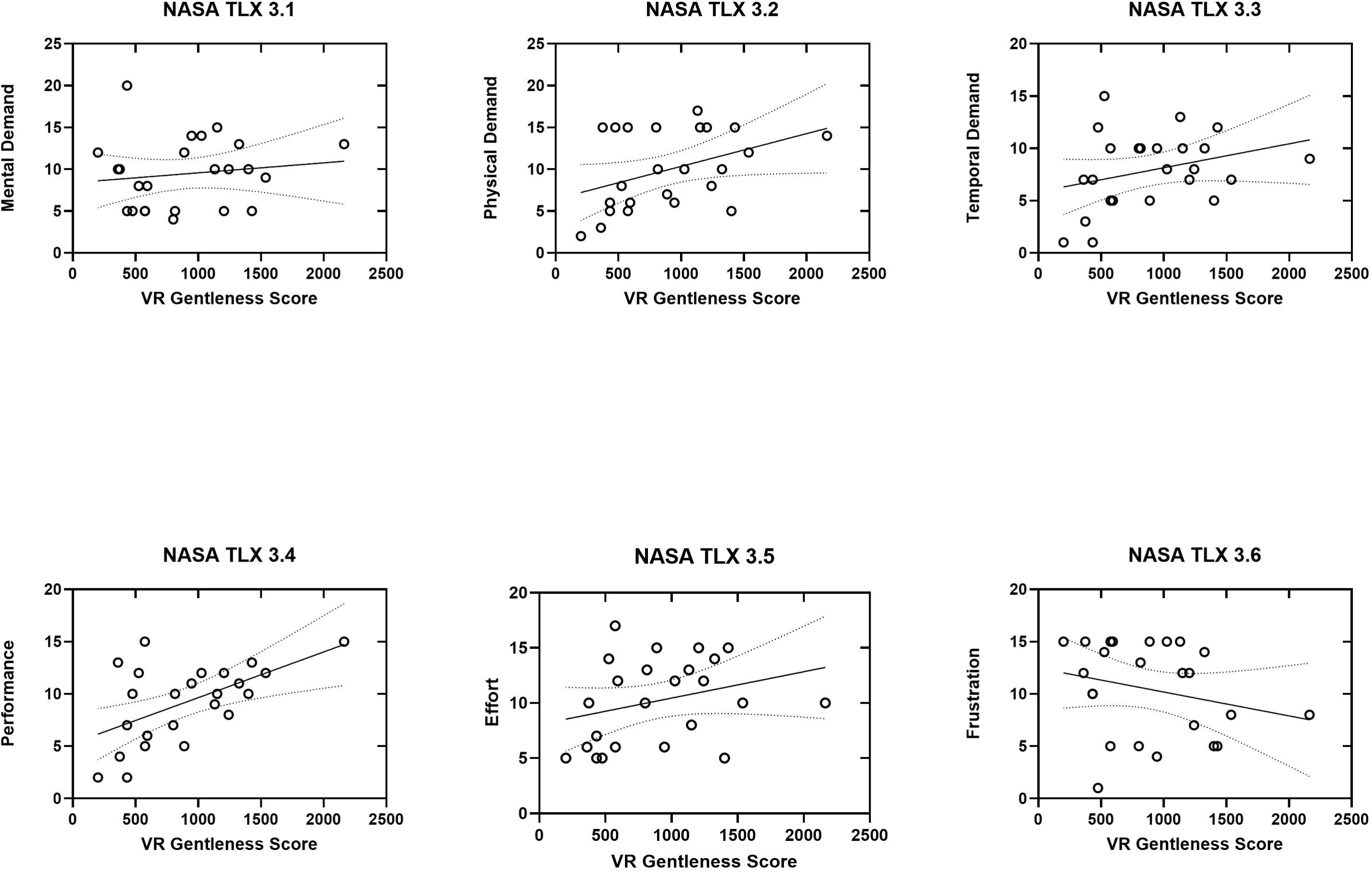
Relationship between VR Gentleness score and NASA-TLX subscales. Scatter plots with linear regression show associations between VR gentleness performance and perceived workload. The VR Gentleness score was positively correlated with Mental Demand (3.1), Physical Demand (3.2), Temporal Demand (3.3), Performance (3.4), and Effort (3.5), whereas Frustration (3.6) showed a slight negative trend. Shaded areas indicate the 95% confidence interval of the regression

For hemodynamic data analysis, we used the concentration change of HbO and HbR for 18 fNIRS channels, with 5 prefrontal (frontal) channels and 4 motor cortex channels in each hemisphere. fNIRS data, which was previously pre-processed by segmenting the continuous signal into non-overlapping 10-second windows and subjected to rigorous artifact rejection (excluding any window whose standard deviation exceeded 4.5 Median Absolute Deviation (MAD) values away from the median), revealed significant and lateralized modulations in cerebral hemodynamic responses based on the Virtual Reality (VR) Score. Each data point (circle) in the Figure 6 represents the aggregated result of a single channel. Specifically, analysis of HbO revealed that in the Right Frontal Cortex, the High VR Score group exhibited a significantly lower concentration of HbO compared to the Low VR Score group (p < 0.05. Conversely, no significant differences were observed for HbO in the Left Frontal Cortex or either motor cortex. A distinct finding emerged in the HbR concentrations: in the Left Motor Cortex, the High VR Score group showed a highly significant increase in HbR concentration compared to the Low VR Score group (p < 0.01). No other significant differences were found for HbR in the right motor or bilateral frontal cortices.

**Figure 6.**
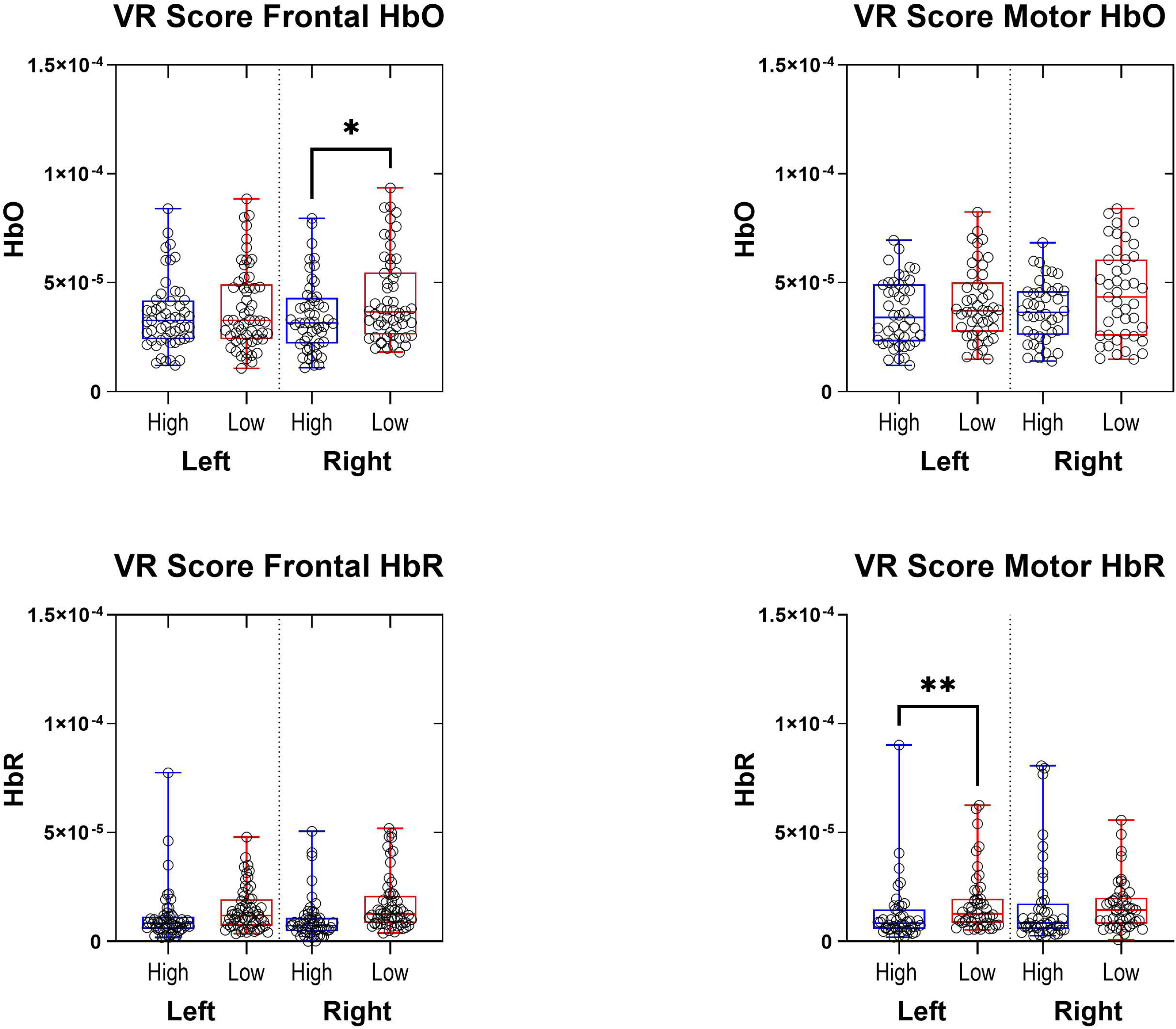
Group-level comparison of frontal and motor cortex hemodynamic responses between high and low VR performance groups. Box plots show oxygenated (HbO) and deoxygenated (HbR) hemoglobin concentrations for left and right frontal and motor regions. Participants were divided into HIGH vs. LOW VR performance groups based on the median VR score.

Table 1 shows that classification performance of fNIRS features in predicting NASA-TLX workload scores was evaluated using three classifiers (RFC, SVC, and KNN) across different feature types (SLOPE, STD, and RMS) and HbO/HbR signals.

For HbO signal, the highest performance was obtained using slope-based features with the Support Vector Classifier (SVC). This configuration achieved an accuracy of 0.8601, F1-score of 0.8718, precision of 0.8395, recall of 0.9067, and an AUC of 0.9071. The HbO Slope + KNN model performed similarly (accuracy = 0.8392, AUC = 0.9134), while the RFC model achieved slightly lower accuracy (0.7692, AUC = 0.8834). Models trained with STD and RMS features of HbO showed moderate classification accuracy, generally between 0.71 and 0.80, with AUC scores below 0.88. Among these, HbO RMS + SVC (accuracy = 0.8042, AUC = 0.8780) and HbO STD + KNN (accuracy = 0.7692, AUC = 0.8508) yielded the best results within their feature categories.

Overall, slope-based HbO features provided the most reliable classification, emphasizing the importance of temporal hemodynamic trends over amplitude variability in predicting gentleness performance. For the deoxyhemoglobin (HbR) signal, classification results were stronger overall compared to HbO. The HbR Slope + RFC and HbR Slope + KNN configurations produced the best performance among all models. Specifically, HbR Slope + RFC reached an accuracy of 0.8881, F1-score of 0.8904, precision of 0.9155, recall of 0.8667, and an AUC of 0.9656, while HbR Slope + KNN achieved comparable accuracy (0.8881) with a slightly lower AUC (0.9325). Within the STD and RMS feature sets, accuracy values remained lower, typically between 0.65 and 0.84. The HbR STD + KNN model performed best among them, with an accuracy of 0.8462 and AUC = 0.9196, while other classifiers (RFC, SVC) yielded weaker results (AUC < 0.80). Overall, HbR slope features consistently outperformed other HbR feature types, exceeding most HbO-based models, confirming that deoxygenated hemoglobin dynamics are more sensitive to variations in cognitive and motor control during VR-based surgical performance.

Based on the normalized confusion matrix results(Figure 7), slope-based models demonstrated the highest overall classification performance across the evaluated machine learning algorithms. The SVC–Slope configuration showed the most balanced classification outcome, with correct prediction rates of 38.5% for the Low VR score group and 47.6% for the High VR score group. RMS- and STD-based feature models resulted in comparatively lower correct classification rates across classifiers, with higher levels of misclassification observed particularly in the Low VR score category. Across all confusion matrices, High VR score predictions were consistently classified with greater accuracy than Low VR scores.

**Figure 7.**
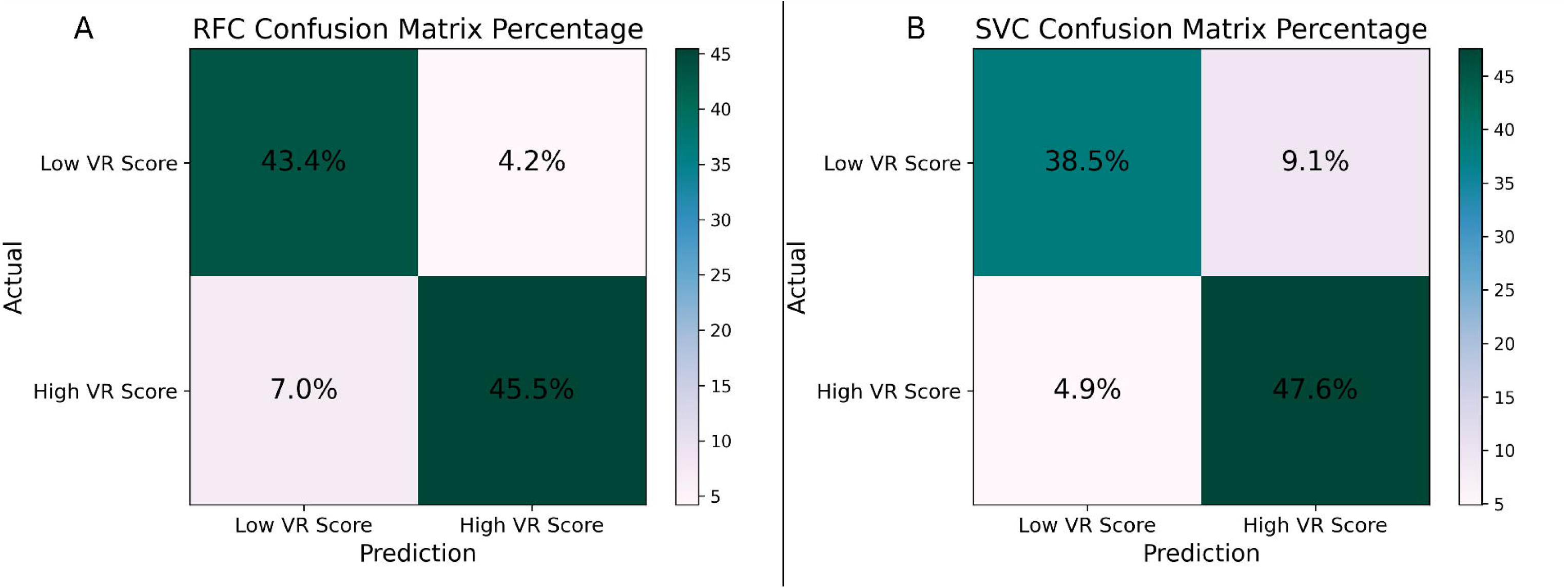
Confusion matrices of the best-performing fNIRS-based models for gentleness classification. **(A)** HbO slope-based features classified using SVC. **(B)** HbR slope-based features classified using RFC.

## DISCUSSION

Our findings provide evidence that gentleness can be assessed using a combination of VR performance metrics and fNIRS-derived cortical signals. While previous VR-based laparoscopic training studies have primarily relied on behavioral and task-based outcomes alone, the present study extends this work by incorporating neural measures of cognitive effort and motor control. To our knowledge, this is the first study to classify gentleness in a simulated laparoscopic task using machine learning applied to hemodynamic features, and to directly compare feature types and model performance within the same framework. These results offer new insight into how cortical activity patterns relate to subtle variations in motor strategy during VR-based surgical training.

The significant relationships observed between the VR Gentleness Score and NASA-TLX subscales indicate a close link between objective motor control and subjective workload perception during VR-based surgical simulation. Specifically, higher gentleness scores were associated with lower physical strain and improved perceived performance, suggesting that participants who applied smoother and more controlled tool forces experienced reduced physical demand and greater task efficiency. The significant association with the total NASA-TLX score further supports the notion that refined psychomotor control contributes to a lower overall workload. These findings align with previous studies demonstrating that expert persons exert lower and more consistent tool–tissue interaction forces, reflecting both improved motor coordination and cognitive regulation during task execution. The integration of the gentleness metric with subjective workload measures thus provides complementary insights into the interplay between cognitive effort and motor precision, reinforcing the potential of force-derived indices as valid indicators of surgical expertise and training progress in immersive VR environments.

The hemodynamic patterns observed in this study provide important insight into the neural processes underlying gentle tissue manipulation during VR-based laparoscopic simulation. The increased HbO and HbR responses observed in the low-performance group, particularly within the right frontal cortex and left motor cortex, suggest a reliance on greater cognitive control and motor correction mechanisms during task execution. This aligns with established evidence showing that early skill acquisition is associated with heightened activation in frontal executive regions as learners engage working memory, error monitoring, response inhibition, and attentional control to maintain task demands. In contrast, participants classified with high gentleness scores demonstrated reduced cortical activation in both frontal and motor regions, suggesting more efficient neural resource allocation and greater procedural fluency. Although both groups were having no prior experience to the laparoscopy or VR gaming, the high VR score group exhibited neural patterns and classification results more typical of advanced learners. This discrepancy may be explained by inter-individual differences in baseline psychomotor aptitude(40) or prior experience with video gaming(41). It is documented that individuals with extensive video gaming experience demonstrate superior visuospatial mapping and faster transition to ’procedural automaticity(42–46). Consequently, the high-gentleness group might have utilized pre-existing neural pathways for spatial navigation and fine motor control, allowing for more efficient neural resource allocation from the outset(47,48).

The significant lateralization effects observed in the left motor cortex further support the interpretation that task proficiency was associated with more automatic motor control. The left hemisphere is known to be dominant in bimanual coordination and sensorimotor integration, particularly in tasks requiring precision and stability, and reduced activation in this region has previously been linked with expert-level performance(32)(49). Similar reductions in prefrontal engagement have been reported in experienced surgeons and skilled tool users, reflecting a progression from deliberate control toward procedural automation as task demands become internalized(50,51). The current findings therefore extend this neuromotor efficiency framework to VR-based assessment of gentleness, demonstrating that subtle behavioral differences in force modulation are mirrored by measurable changes in cortical activation.

Together, these results suggest that fNIRS-derived hemodynamic signals may serve as meaningful biomarkers of surgical gentleness, capturing differences in motor strategy that may not be evident from behavioral metrics alone. The observed relationship between reduced prefrontal engagement and gentler manipulation supports the potential use of neuroadaptive feedback systems to guide learners toward more efficient and safer instrument handling patterns during simulation-based training.

The fNIRS-based classification of the VR Gentleness Score revealed that slope features derived from HbO and HbR signals provided the highest discriminative performance across classifiers. Among all models, HbR Slope + RFC and HbR Slope + KNN achieved the best accuracy values (0.8881) with AUC scores of 0.9656 and 0.9325, respectively, while the best-performing HbO model (HbO Slope + SVC) reached 0.8601 accuracy and AUC = 0.9071. These findings indicate that the temporal evolution of both oxy- and deoxyhemoglobin concentrations is more informative than static or variability-based features (STD, RMS) for differentiating levels of motor control and movement smoothness during VR-based surgical tasks.

The superior performance of HbR features compared to HbO aligns with evidence that deoxygenated hemoglobin changes more directly reflect localized neuronal activation and are less affected by systemic physiological artifacts (52). The slope parameter captures the rate of task-related cortical activation, offering a sensitive index of how effectively participants regulate force and precision while interacting with the virtual environment(53).

From a methodological perspective, the superior performance of KNN may be attributed to its non-parametric nature and ability to model non-linear relationships between hemodynamic patterns and subjective workload levels. While more complex algorithms (e.g., RFC, SVC) performed comparably on certain feature sets, KNN consistently provided the most balanced precision–recall profile, suggesting robustness for small-sample datasets.

Integrating these findings with behavioral metrics, such as the VR Gentleness Score, underscores the complementary nature of neurophysiological and psychomotor indicators in assessing cognitive–motor efficiency. Participants demonstrating smoother and more controlled tool-force modulation also exhibited lower subjective workload and stronger HbR slope responses, supporting the notion that optimized motor control is accompanied by efficient cortical resource allocation. Together, these outcomes highlight the potential of multimodal frameworks, combining VR performance data and fNIRS features for objective and adaptive assessment of surgical training performance.

Overall, these results suggest that HbR slope dynamics serve as a robust neurophysiological marker of fine motor performance, while HbO slope responses provide complementary information about overall cortical oxygenation during task execution. Together, they support the utility of fNIRS in capturing performance-related cortical changes associated with the VR Gentleness Score.

This study has several limitations that should be considered when interpreting the findings. First, the number of participants was relatively small, which limits the statistical power of the analyses and may have constrained the performance and generalizability of the machine learning models. A larger sample size would allow more robust model training and validation, increase confidence in classification outcomes, and potentially reveal subtler neurobehavioral differences related to skill level. Furthermore, a comprehensive assessment of baseline motor aptitude or visuomotor proficiency was not conducted. The haptic feedback provided by the VR simulator was limited and did not fully replicate the tactile characteristics of real laparoscopic tissue interaction, which may have influenced gentleness-related behavioral and neural responses.

## CONCLUSION

This study demonstrates that gentleness during VR-based laparoscopic simulation can be quantified using fNIRS-derived hemodynamic features and machine learning classification. Task-related hemodynamic activity in frontal and motor regions differed between participants with high and low VR gentleness scores, indicating that fNIRS measures are sensitive to performance-related neural differences during laparoscopic tasks performed in a virtual reality environment. The successful classification of gentleness using slope-based features highlights the potential of incorporating temporal neural metrics into automated performance assessment frameworks. As VR training platforms continue to evolve, the integration of real-time neurophysiological monitoring may support the development of adaptive feedback systems capable of promoting more efficient motor strategies and safer surgical behaviors. Future work with larger and more diverse samples including experienced participant (e.g. surgeons), baseline motor aptitude testing, higher-fidelity haptics, and longitudinal study designs will be essential for refining these methods and evaluating their applicability in real-world surgical education.

## ACKNOWLEDGEMENTS

The authors thank Prof. Doğa Demirel for the design and development of the virtual reality–based Laparoscopic Double Grasper Task used in this study. We would like to thank Cagri Zengin for the support of machine learning analysis of this manuscript.

